# PP2A^B55δ^ Responsible for the High Initial Rates of Alcoholic Fermentation in Sake Yeast Strains of *Saccharomyces cerevisiae*

**DOI:** 10.1101/402081

**Authors:** Daisuke Watanabe, Takuma Kajihara, Yukiko Sugimoto, Kenichi Takagi, Megumi Mizuno, Yan Zhou, Jiawen Chen, Kojiro Takeda, Hisashi Tatebe, Kazuhiro Shiozaki, Nobushige Nakazawa, Shingo Izawa, Takeshi Akao, Hitoshi Shimoi, Tatsuya Maeda, Hiroshi Takagi

## Abstract

Sake yeast strain Kyokai no. 7 (K7) and its *Saccharomyces cerevisiae* relatives carry a homozygous loss-of-function mutation in the *RIM15* gene, which encodes a Greatwall-family protein kinase. Disruption of *RIM15* in non-sake yeast strains leads to improved alcoholic fermentation, indicating that the defect in Rim15p is associated with the enhanced fermentation performance of sake yeast cells. In order to understand how Rim15p mediates fermentation control, we here focused on target-of-rapamycin protein kinase complex 1 (TORC1) and protein phosphatase 2A with the B55Δ regulatory subunit (PP2A^B55δ^), complexes that are known to act upstream and downstream of Rim15p, respectively. Several lines of evidence, including our previous transcriptomic analysis data, suggested enhanced TORC1 signaling in sake yeast cells during sake fermentation. Fermentation tests of the TORC1-related mutants using a laboratory strain revealed that TORC1 signaling positively regulates the initial fermentation rate in a Rim15p-dependent manner. Deletion of the *CDC55* gene encoding B55δ abolished the high fermentation performance of Rim15p-deficient laboratory yeast and sake yeast cells, indicating that PP2A^B55δ^ mediates the fermentation control by TORC1 and Rim15p. The TORC1-Greatwall-PP2A^B55δ^ pathway similarly affected the fermentation rate in the fission yeast *Schizosaccharomyces pombe*, strongly suggested that the evolutionarily conserved pathway governs alcoholic fermentation in yeasts. It is likely that elevated PP2A^B55δ^ activity accounts for the high fermentation performance of sake yeast cells. Heterozygous loss-of-function mutations in *CDC55* found in K7-related sake strains may indicate that the Rim15p-deficient phenotypes are disadvantageous to cell survival.

**IMPORTANCE:** The biochemical processes and enzymes responsible for glycolysis and alcoholic fermentation by the yeast *S. cerevisiae* have long been the subject of scientific research. Nevertheless, the factors determining fermentation performance *in vivo* are not fully understood. As a result, the industrial breeding of yeast strains has required empirical characterization of fermentation by screening numerous mutants through laborious fermentation tests. To establish a rational and efficient breeding strategy, key regulators of alcoholic fermentation need to be identified. In the present study, we focused on how sake yeast strains of *S. cerevisiae* have acquired high alcoholic fermentation performance. Our findings provide a rational molecular basis to design yeast strains with optimal fermentation performance for production of alcoholic beverages and bioethanol. In addition, as the evolutionarily conserved TORC1-Greatwall-PP2A^B55δ^ pathway plays a major role in the glycolytic control, our work may contribute to research on carbohydrate metabolism in higher eukaryotes.

## INTRODUCTION

Sake, an alcoholic beverage made from fermented rice, typically has a higher alcohol content than beer or wine. During sake fermentation, saccharification by hydrolytic enzymes of *Aspergillus oryzae* and alcoholic fermentation by *Saccharomyces cerevisiae* sake yeast are the major bioconversions. Thus, the high alcohol content of sake is at least partly attributable to the unique characteristics of sake yeast. Sake yeast strains have long been selected based on the high fermentation performance, as well as the balanced production of aroma and flavor compounds (1, 2). Our previous comparative genomic and transcriptomic analyses revealed that a representative sake yeast, strain Kyokai no. 7 (K7), and its relatives carry a loss-of-function mutation in *RIM15* (*rim15*^*5054_5055insA*^), a gene of a highly conserved Greatwall-family protein kinase (3–5). Disruption of the *RIM15* gene in non-sake yeast strains, such as laboratory, beer, and bioethanol strains, leads to an increase in the fermentation rate (5–9), demonstrating that Rim15p inhibits alcoholic fermentation. Thus, the *rim15*^*5054_5055insA*^ mutation appears to be associated with the enhanced fermentation property of K7. Nevertheless, this loss-of-function mutation cannot be solely responsible for the sake yeast’s improved fermentation, because expression of the functional *RIM15* gene does not suppress alcoholic fermentation in K7 (5). To better understand this phenomenon, the Rim15p-mediated fermentation control needs to be further dissected through comparative analysis between sake and non-sake yeast strains.

While Rim15p has been identified as a key inhibitor of alcoholic fermentation, involvement of the upstream regulators of Rim15p (Fig. S1) in fermentation control has not yet been fully examined. In *S. cerevisiae*, Rim15p activity is under the control of several nutrient-sensing signaling protein kinases, including protein kinase A (PKA), the phosphate-sensing cyclin and cyclin-dependent protein kinase (CDK) complex termed Pho80p-Pho85p, and target-of-rapamycin protein kinase complex 1 (TORC1) (10, 11). Thus, inactivation of these kinases under nutrient starvation or other stress conditions may trigger Rim15p-dependent inhibition of alcoholic fermentation. Activation of TORC1 is mediated by the heterodimeric Rag GTPases (Gtr1p-Gtr2p in *S. cerevisiae*), which are negatively regulated by the Seh1p-associated protein complex inhibiting TORC1 (SEACIT) subcomplex (Iml1p-Npr2p-Npr3p in *S. cerevisiae*) that acts as a GTPase-activating protein for Gtr1p (12–14). Active TORC1 phosphorylates multiple targets including Sch9p, the yeast orthologue of the mammalian serum and glucocorticoid-regulated kinase (SGK) (15). Direct phosphorylation of Rim15p at Ser1061 by Sch9p contributes to sequestration of Rim15p in the cytoplasm, thereby inhibiting Rim15p functions (16). Recently, it was reported that the TORC1-Sch9p-Rim15p pathway is conserved and present in the evolutionarily distant fission yeast *Schizosaccharomyces pombe* (17), although it remains to be determined if the pathway affects the fermentation performance in this yeast species. In contrast, mammalian TORC1 (mTORC1) positively regulates glycolysis by the induction of glycolytic gene expression through hypoxia-inducible factor 1α (HIF1α) (18).

In *S. cerevisiae*, Rim15p targets the redundant transcription factors Msn2p and Msn4p (Msn2/4p) to mediate entry into the quiescent state (19, 20). In the context of fermentation control, Rim15p and Msn2/4p are required for the transcriptional induction of the UDP-glucose pyrophosphorylase-encoding gene *UGP1*, which switches the mode of glucose metabolism from glycolysis (a catabolic mode) to UDP-glucose synthesis (an anabolic mode) (7). However, no orthologue of Msn2/4p has been found in other organisms, and the role of the Greatwall-family protein kinases in carbohydrate metabolism is unknown in *S. pombe* or higher eukaryotes. The Greatwall protein kinase was originally identified as a potential cell cycle activator in *Drosophila* (21). In animals, Greatwall directly phosphorylates a small protein called α-endosulfine (ENSA), which inhibits the activity of protein phosphatase 2A accompanied by a regulatory subunit B55Δ (PP2A^B55δ^) (22, 23). Due to the antimitotic activity of PP2A^B55δ^, Greatwall is required for maintenance of mitosis. More recently, the Greatwall-ENSA-PP2A^B55δ^ pathway was reported to be conserved in *S. cerevisiae*; Rim15p phosphorylates ENSA orthologues Igo1/2p to inhibit PP2A with the Cdc55p regulatory subunit (24–26). The orthologous pathway has also been found in *S. pombe* and it plays a pivotal role in TORC1-mediated cell cycle control (17). However, to our knowledge, the effect of PP2A^B55δ^ on fermentation performance has not previously been described.

In the present study, we tested whether the TORC1-Greatwall-PP2A^B55δ^ pathway participates in the control of alcoholic fermentation in *S. cerevisiae* and *S. pombe*. Our results provide new insights into how yeast cells determine the mode of glucose metabolism, especially in the context of the enhanced fermentation performance of sake yeast strains.

## RESULTS

### TORC1-associated transcriptomic profiles during alcoholic fermentation in laboratory and sake yeast strains

Our previous comparative transcriptomic analysis indicated that the expression of the Rim15p- and Msn2/4p-targeted genes was attenuated in K701 (a strain derived from K7) compared to that in the laboratory strain X2180 early in a 20-d sake fermentation test (3). This may be attributed not only to the sake yeast-specific loss-of-function mutation in the *RIM15* gene (*rim15*^*5054_5055insA*^; see also ref. 5), but also to higher TORC1 activity in the sake strains. TORC1 activity induces the ribosomal genes and the ribosome biogenesis genes, while it represses the nitrogen catabolite repression (NCR) and general amino acid control (GAAC) genes, as well as the Rim15p- and Msn2/4p-dependent stress-response genes (27) (Fig. S1). We found that these transcriptomic traits under the control of TORC1 are coordinated in common during sake fermentation. Although the Sfp1p-targeted genes encoding the ribosome-associated proteins in X2180 were rapidly downregulated during the progression of alcoholic fermentation, K701 cells maintained higher levels of these mRNAs at the initial stage of fermentation (Fig. 1A). Whereas the NCR and GAAC genes in X2180 were transiently upregulated early in sake fermentation, this transcriptional induction was not observed in K701 (Fig. 1B). Together, these data suggested that inactivation of TORC1 after the onset of alcoholic fermentation leads to attenuation of the Sfp1p-targeted gene expression and induction of the NCR and GAAC regulons in X2180. This phenomenon was less clearly observed in K701, implying a slower decline in TORC1 activity during the initial stage of alcoholic fermentation by K701.

**FIG 1.**
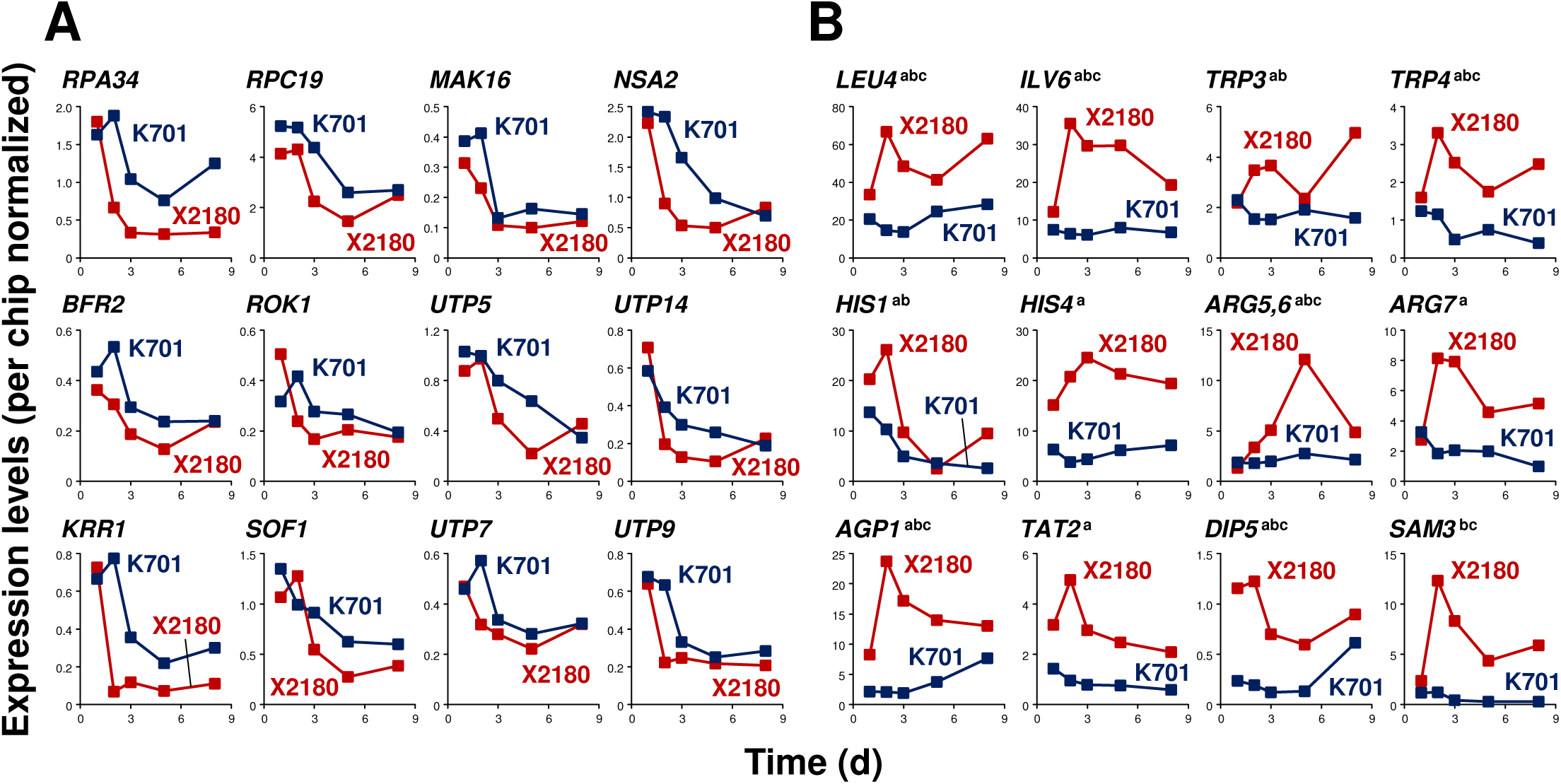
Gene expression profiles downstream of TORC1: Comparison between a laboratory strain and a sake strain during sake fermentation. (A) Expression profiles of the ribosome-associated genes under the control of Sfp1p. (B) Expression profiles of genes belonging to the NCR and GAAC regulons. Expression levels in a laboratory strain (X2180) and a sake strain (K701) are derived from our previous DNA microarray data (3) and are indicated by red and blue, respectively. TORC1, target-of-rapamycin complex 1; NCR, nitrogen catabolite repression; GAAC, general amino acid control.

To directly monitor TORC1 activity, an antibody against phospho-Thr737 of Sch9p (28) was used, as this TORC1-dependent phosphorylation of Sch9p is known to mediate signaling to Rim15p. Laboratory yeast and K701-lineage sake yeast cells engineered to overexpress 3HA-tagged Sch9p from a glycolytic gene promoter were sampled during alcoholic fermentation in YPD20 medium. The sake strain exhibited a higher rate of carbon dioxide emission and completed alcoholic fermentation more rapidly than the laboratory strain (Fig. S2A). Phosphorylation of Sch9p Thr737 was detected only at the initial stage (at 6 h from the onset of alcoholic fermentation), and was more prominent in the sake strain than in the laboratory strain (Fig. S2B). The signals decayed quickly over time in both strains, suggesting that TORC1 activity is highest at the onset of alcoholic fermentation. It should be noted that the glycolytic promoter used in this study was inactivated after the completion of logarithmic phase (> 12 - 24 h) and 3HA-Sch9p was expressed only at low levels after 2 days.

### Effects of the TORC1-Greatwall-PP2A^B55δ^ pathway on fermentation performance

In *S. cerevisiae* laboratory strains, loss of Rim15p leads to an increase in the initial rate of carbon dioxide emission during alcoholic fermentation (5, 7) (Fig. 2A). To examine whether TORC1 acts as a negative regulator of Rim15p activity in fermentation control, we tested the effect of altered TORC1 signaling on fermentation performance in a laboratory strain. Addition of a low concentration (1 nM) of the TORC1 inhibitor rapamycin to the medium led to a decrease in the rate of carbon dioxide emission from 1.5 d to 4 d (Fig. 2B). Since cell growth was not severely affected by 1 nM rapamycin (data not shown), the observed attenuation of carbon dioxide production was most likely indicative of reduced cellular fermentation performance. Deletion of the *TOR1* gene, which encodes a nonessential catalytic subunit of TORC1, also decreased carbon dioxide emission from 1.5 d to 3.5 d (Fig. 2C). Deletion of *TOR2*, which encodes a second TOR kinase that can also serve as a catalytic subunit of TORC1, was not tested in this study because Tor2p is essential for cell viability in *S. cerevisiae*. In contrast, the hyperactive *TOR1* and *TOR2* alleles (*TOR1*^*L2134M*^ and *TOR2*^*L2138M*^) (28) increased carbon dioxide emission around 1 d to 2 d (Figs. 2D and E). Strains harboring either of these hyperactive alleles exhibited drastic decreases in the rate of carbon dioxide emission toward the end of the fermentation tests. These results suggested that TORC1 activity correlates with fermentation performance during the initial stage of the process. Indeed, deletion of *GTR1* or *GTR2*, activators of TORC1 signaling, decreased carbon dioxide emission (Figs. 2F and G). In addition, disruption of *NPR2* or *NPR3*, which encode the components of the SEACIT subcomplex that inhibit TORC1 signaling, resulted in increased carbon dioxide emission around 1.5 d to 2 d (Figs. 2H and I), corroborating the role of TORC1 as a positive regulator of alcoholic fermentation. On the other hand, loss of Sch9p, which mediates signaling between TORC1 and Rim15p (Fig. S1), markedly decreased carbon dioxide emission (Fig. 2J).

**FIG 2.**
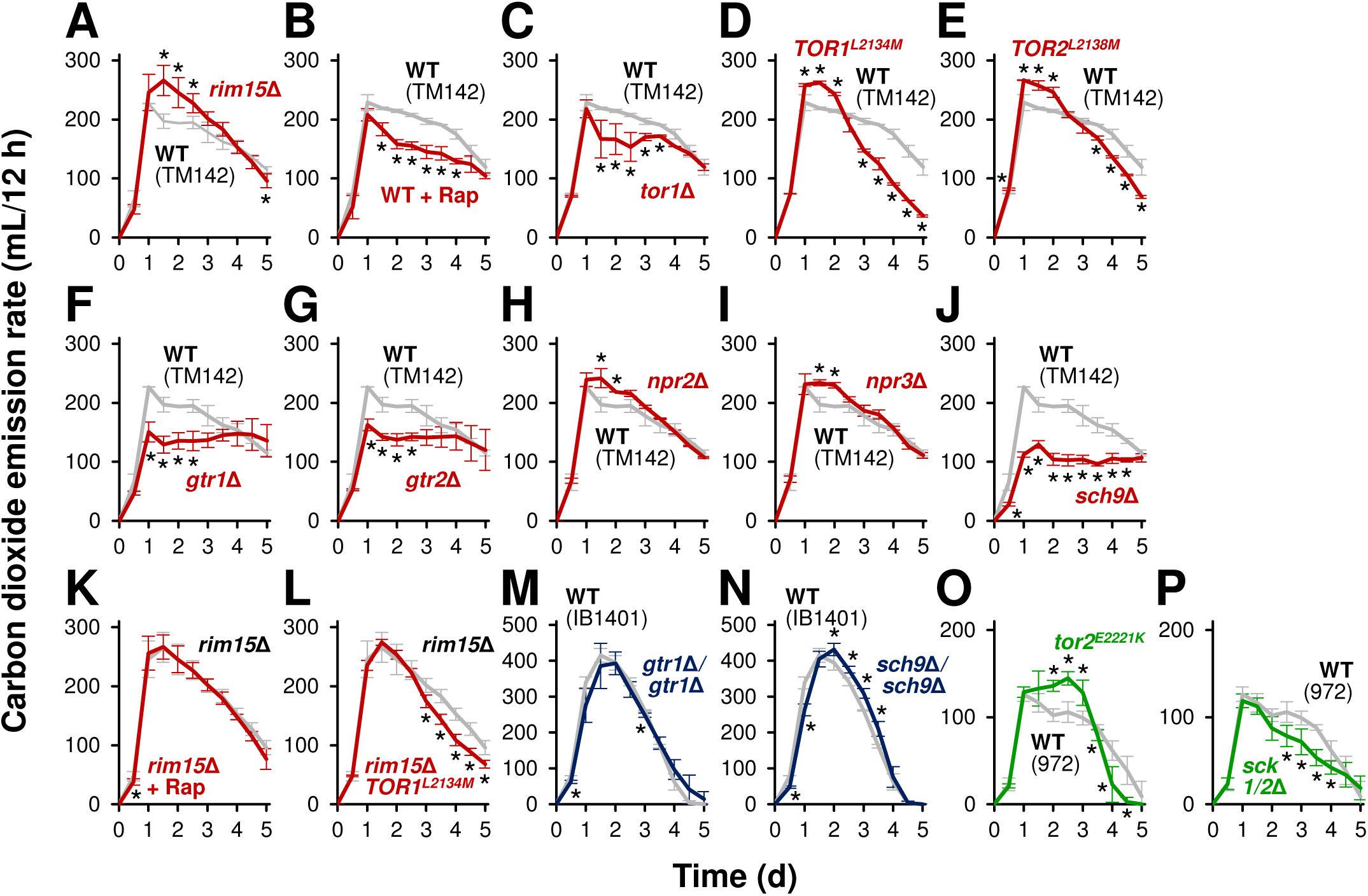
Effects of modification of the TORC1-Greatwall pathway on fermentation progression. Fermentation was monitored by measuring carbon dioxide emission. (A) Fermentation profiles of strain TM142 (wild type; gray) and its *rim15*Δ disruptant (red). (B) Fermentation profiles of strain TM142 in YPD20 medium in the absence (wild type, gray) or presence (red) of 1 nM rapamycin. (C to J) Fermentation profiles of strain TM142 (wild type; gray) and its *tor1*Δ (C), *TOR1*^*L2134M*^ (D), *TOR2*^*L2138M*^ (E), *gtr1*Δ (F), *gtr2*Δ (G), *npr2*Δ (H), *npr3*Δ (I), or *sch9Δ* (J) mutant (red). (K) Fermentation profiles of strain TM142 *rim15*Δ in YPD20 medium in the absence (*rim15*Δ; gray) or presence (red) of 1 nM rapamycin. (L) Fermentation profiles of strain TM142 *rim15*Δ (*rim15*Δ; gray) and its *TOR1*^*L2134M*^ mutant (red). (M, N) Fermentation profiles of strain IB1401 (wild type; gray) and its *gtr1*Δ*/gtr1*Δ(M) or *sch9Δ/sch9Δ* (N) disruptant (blue). (O, P) Fermentation profiles of the wild-type *S. pombe* strain (wild type; gray) and its *tor2*^*E2221K*^ (O) or *sck1/2*Δ (P) mutant (green). Fermentation tests were performed in YPD20 medium (A to N) or in YPD10 medium (O, P) at 30°C for 5 d. Values represent the mean ± SD of data from two or more independent experiments. *, significantly different from the value for the control experiment (*t* test, *P* < 0.05). Note that the experiments using laboratory, sake, and fission yeast strains are indicated in red, blue, and green, respectively. WT, wild type; Rap, rapamycin.

Next, the effects of TORC1 on fermentation performance were examined in the Rim15p-deficient strains. In *rim15*Δ cells of the laboratory strain, 1 nM rapamycin did not affect carbon dioxide emission (Fig. 2K). We confirmed that the growth of *rim15*Δ cells was not affected by 1 nM rapamycin (data not shown). In the *rim15*Δ background, the hyperactive *TOR1*^*L2134M*^ allele did not increase the initial rate of carbon dioxide emission (1 - 2 d), but caused a decrease in the later stage of fermentation (> 3 d) (Fig. 2L). These data suggested that Rim15p is required for the TORC1-triggered fermentation control, specifically in the early stage of alcoholic fermentation. Thus, it was predicted that the fermentation performance of the sake strain defective in Rim15p is not affected by TORC1 signaling. As expected, deletion of the *GTR1* or *SCH9* genes in the sake strain did not change the maximum rate of carbon dioxide emission (Figs. 2M and N), although alcoholic fermentation was slightly delayed in both cases, probably due to slower cell growth. We also assessed whether the conserved TORC1-Greatwall pathway affects fermentation performance in the fission yeast *S. pombe*. As observed in budding yeast, an activated allele of *tor2*^*+*^, *tor2*^*E2221K*^ (29), brought about increased carbon dioxide emission in fission yeast (Fig. 2O). Furthermore, deletion of the redundant Sch9p-orthologous genes, *sck1*^*+*^ and *sck2*^*+*^, resulted in decreased carbon dioxide emission (Fig. 2P).

Does Greatwall-triggered signaling to PP2A^B55δ^ play a role in the control of alcoholic fermentation? Deletion of the redundant ENSA-encoding genes *IGO1* and *IGO2* (*IGO1/2*), which mediate the signaling between Greatwall and PP2A^B55δ^ *in S. cerevisiae* (Fig. S1), led to an increased rate of carbon dioxide emission, as did deletion of *RIM15* (Figs. 3A and B). Similarly, in *S. pombe*, both the *cek1*Δ *ppk18*Δ and *igo1*Δ strains, which lack Greatwall and ENSA, respectively (17), exhibited higher rates of carbon dioxide emission than did the wild type (Figs. 3C and D). PP2A is a heterotrimeric enzyme complex composed of structural (A), regulatory (B), and catalytic (C) subunits. In budding yeast, the loss of both C subunit-encoding genes *PPH21* and *PPH22* leads to cell death, but disruption of either gene alone only weakly decreased carbon dioxide emission (Figs. 3E and F). In addition, deletion of the A subunit-encoding *TPD3* gene inhibited alcoholic fermentation (Fig. 3G). Moreover, deletion of *CDC55*, which encodes a B55Δ-family regulatory subunit, severely decreased the rate of carbon dioxide emission throughout the duration of fermentation, whereas deletion of *RTS1* that encodes a B56-family regulatory subunit promoted alcoholic fermentation (Figs. 3H and I). In *S. pombe*, loss of the *ppa1*^*+*^ or *ppa2*^*+*^ gene, which encode C subunit isoforms, did not appear to affect alcoholic fermentation; however, loss of the *pab1*^*+*^ gene encoding a B55Δ subunit impaired alcoholic fermentation (Figs. 3J–L). Together, these data suggested that the Greatwall-ENSA-PP2A^B55δ^ pathway is involved in the control of alcoholic fermentation in both *S. cerevisiae* and *S. pombe*. When combined with the Greatwall or ENSA defects, deletion of the *CDC55* (*S. cerevisiae*) or *pab1*^*+*^ (*S. pombe*) genes almost fully canceled high fermentation performance (Figs. 3M–O). Thus, PP2A^B55δ^ is likely the major target of Greatwall and ENSA in the control of alcoholic fermentation in both yeasts.

**FIG 3.**
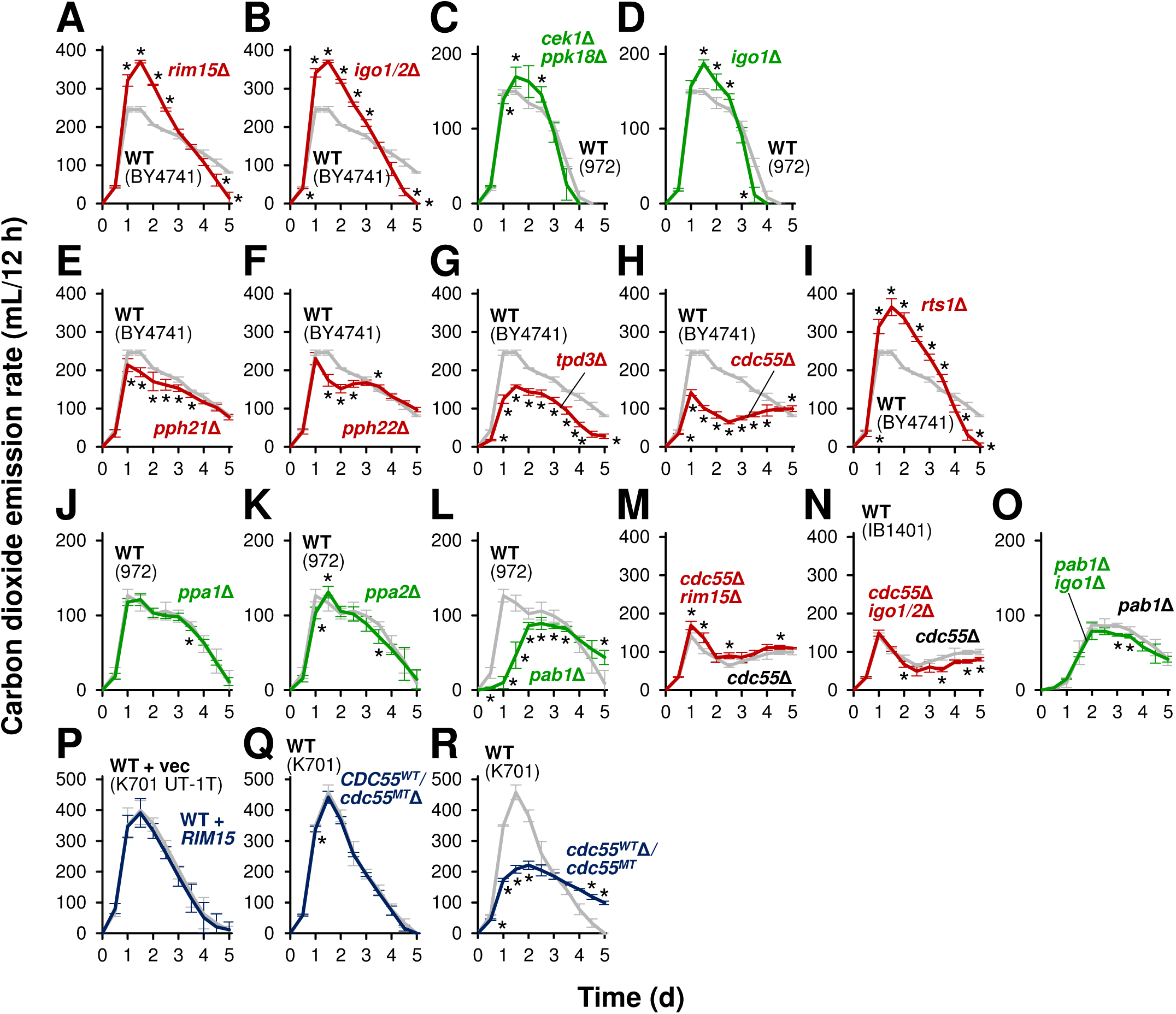
Effects of modification of the Greatwall-PP2A^B55Δ^ pathway on fermentation progression. Fermentation was monitored by measuring carbon dioxide emission. (A, B) Fermentation profiles of strain BY4741 (wild type; gray) and its *rim15*Δ (A) or *igo1/2*Δ (B) disruptant (red). (C, D) Fermentation profiles of the wild-type *S. pombe* strain (wild type; gray) and its *cek1*Δ*/ppk18*Δ (C) or *igo1*Δ (D) disruptant (green). (E to I) Fermentation profiles of strain BY4741 (wild type; gray) and its *pph21*Δ (E), *pph22*Δ (F), *tpd3*Δ (G), *cdc55*Δ (H), or *rts1*Δ (I) disruptant (red). (J to L) Fermentation profiles of the wild-type *S. pombe* strain (wild type; gray) and its *ppa1*Δ (J), *ppa2*Δ (K) or *pab1*Δ (L) disruptant (green). (M, N) Fermentation profiles of strain BY4741 *cdc55*Δ (*cdc55*Δ; gray) and its *rim15*Δ (M) or *igo1/2*Δ (N) disruptant (red). (O) Fermentation profiles of the *S. pombe pab1*Δ strain (*pab1*Δ; gray) and its *igo1*Δ disruptant (green). (P) Fermentation profiles of strain K701 UT-1T with an empty vector (wild type; gray) and with a functional *RIM15*-expressing plasmid (blue). (Q, R) Fermentation profiles of strain K701 (wild type; gray) and its *CDC55*^*WT*^*/cdc55*^*MT*^δ (Q) or *cdc55*^*WT*^δ*/cdc55*^*MT*^ (N) disruptant (blue). Fermentation tests were performed in YPD20 medium (A, B, E to I, M, N, P to R) or in YPD10 medium (C, D, J to L, O) at 30°C for 5 d. Values represent the mean ± SD of data from two or more independent experiments. *, significantly different from the value for the control experiment (*t* test, *P* < 0.05). Note that the experiments using laboratory, sake, and fission yeast strains are indicated by red, blue, and green, respectively. WT, wild type.

Consistent with a previous report (5), expression of the functional *RIM15* gene derived from a laboratory strain did not attenuate alcoholic fermentation in the sake strain (Fig. 3P). Therefore, we next evaluated the role of PP2A^B55δ^ downstream of Rim15p in the high fermentation performance of sake yeast cells. Interestingly, we found that the diploid sake strain K701 is heterozygous for the deletion of a single adenine nucleotide at position 1092 of the *CDC55* gene (designated the *cdc55*^*MT*^ allele), resulting in a frameshift and premature polypeptide termination (Fig. 4A); thus, K701 carries only one functional *CDC55* allele (designated the *CDC55*^*WT*^ allele). To directly test the role of PP2A^B55δ^ in the sake yeast, the K701 strain was mutagenized by introduction of a *CDC55*-disrupting construct, yielding 12 heterozygous disruptants. Direct sequencing of the *CDC55* loci amplified from genomic DNA revealed that the *cdc55*^*MT*^ allele was disrupted in six of the heterozygous disruptants, while the *CDC55*^*WT*^ allele was disrupted in the other six heterozygous disruptants. The former class, in which the *CDC55*^*WT*^ allele remains intact, exhibited fermentation characteristics similar to the parental K701 strain (Fig. 3Q), while the latter class with no functional *CDC55* gene exhibited markedly lower carbon dioxide emission, especially in the initial stage of fermentation (0.5 - 2 d) (Fig. 3R). These results indicated that the *CDC55*^*WT*^ allele is required for the high fermentation performance of the K701 sake yeast strain.

**FIG 4.**
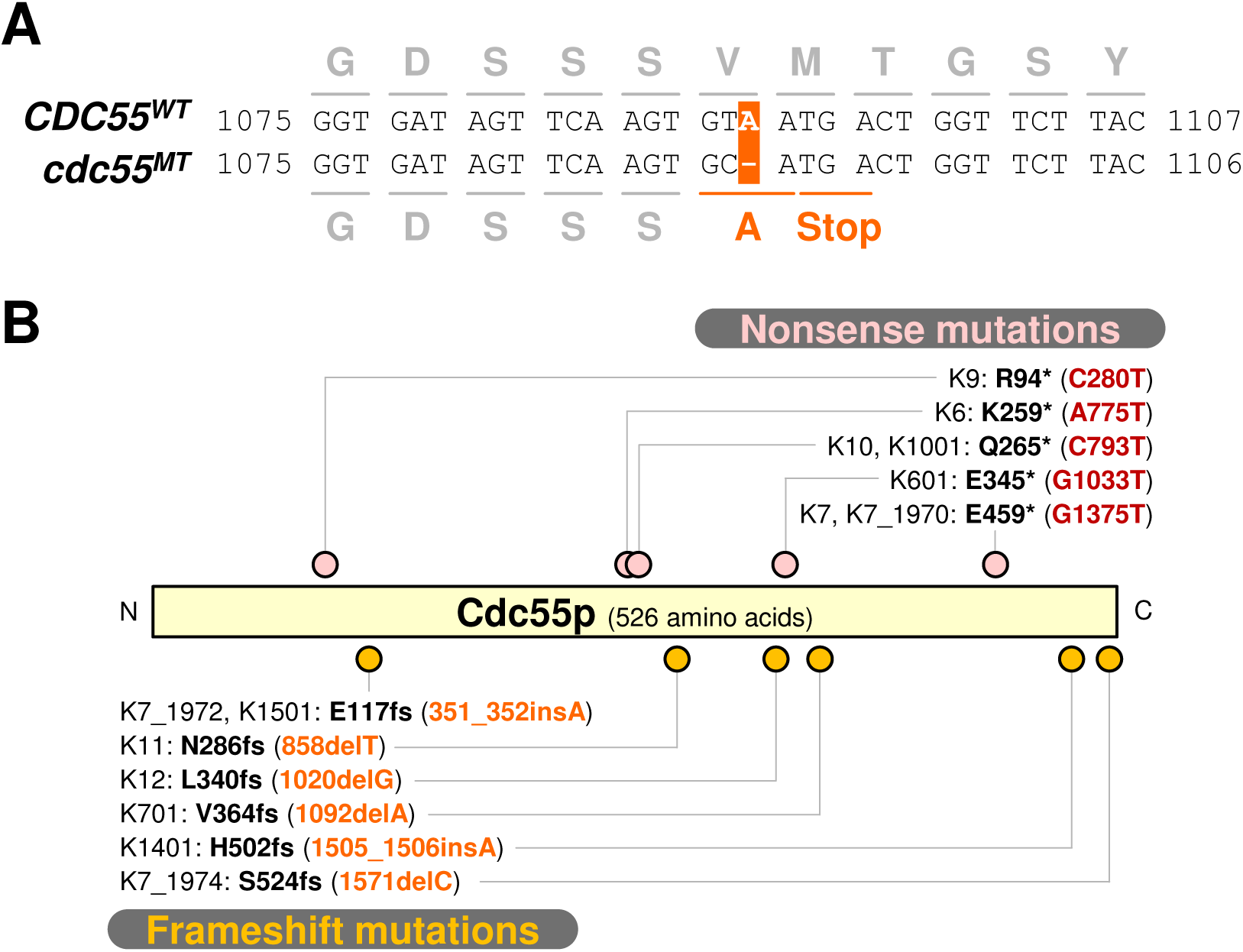
Heterozygous nonsense or frameshift mutations found in the *CDC55* genes of K7-related sake strains. (A) The *cdc55*^*1092delA*^ (a.k.a. *cdc55*^*MT*^) mutation unique to K701. In this loss-of-function allele of K701, deletion of a single adenine nucleotide at ORF nucleotide 1092 causes a premature stop codon. (B) Mutation sites of the *CDC55* gene of K7-related sake strains. Nonsense and frameshift mutation sites are indicated by pink and orange dots, respectively. fs, frameshift.

### Heterozygous nonsense or frameshift mutations in the *CDC55* gene of the sake yeast strains

As mentioned above, we identified a heterozygous loss-of-function mutation (*cdc55*^*1092delA*^) in diploid K701 (Fig. 4A). To test whether this mutation is conserved among the sake strains, we analyzed the sequence of the *CDC55* genes in 17 K7-related Kyokai sake strains, including K6, K601, K7, K701, K9, K901, K10, K1001, K11, K12, K13, K14, K1401, K1501, K1601, K1701, and K1801 [Note that the numbering corresponds to the sequential isolation of these strains. The “-01” suffix is used to indicate foamless variants that do not generate thick foam layers during sake fermentation; for instance, K701 is the foamless variant of K7 (30).]. As shown in Fig. 4B, the *cdc55*^*1092delA*^ mutation is unique to K701, and 65% (11 of 17) of the tested strains contain other nonsense or frameshift mutations in the open reading frame of the *CDC55* gene. Notably, the three most recently isolated strains, K1601, K1701, and K1801, have neither a nonsense mutation nor a frameshift mutation in this locus. Although there are a few lineage-specific mutations, such as *cdc55*^*C793T*^ in K10 and K1001 and *cdc55*^*351_352insA*^ in K7 and K1501 (2), closely associated strains do not always contain the same mutation (e.g., K6 versus K601, K7 versus K701, or K7 versus K11). Each year, every Kyokai sake yeast strain was selected from clone stocks before distribution by the Brewing Society of Japan; notably, the K7 strains from three different years (1970, 1972, and 1974) carry distinct *cdc55* mutations. The K7 strain used for whole-genome analysis (4) harbors a *cdc55* mutation identical to the *cdc55*^*1571delC*^ allele in K7_1970. Thus, it appears that most of the *cdc55* mutations represent independent events that occurred after the establishment of the individual sake strains. While the *cdc55*^*1571delC*^ mutation in K7_1974 results in additional 27 amino acid residues at the carboxyl terminus of the encoded protein, each of the other frameshift mutations leads to a premature stop codon that truncates the carboxyl terminus. Since all of the identified mutations are heterozygous, the effects of the *cdc55* loss-of-function mutations may be masked by the functional *CDC55* allele, as observed in K701.

### Effects of PP2A^B55δ^ on the intracellular levels of glycolytic intermediates

Since PP2A^B55δ^ dephosphorylates many cellular substrates (31), it is difficult to infer how PP2A^B55δ^ controls alcoholic fermentation. However, it may be worth examining whether PP2A^B55δ^ regulates the activities of carbon metabolic enzymes through protein dephosphorylation as several recent studies have shed light on posttranslational modification as regulatory mechanisms for metabolic flux *in vivo* (32, 33). In the present study, we adopted a metabolomic approach to explore the glycolytic reactions that may be affected by the loss of PP2A^B55δ^ function. Metabolites were extracted from cells sampled at the early stages (6 h, 1 d, or 2 d) of alcoholic fermentation in YPD20 medium. Relative metabolite levels at 6 h indicated that the pools of early glycolytic intermediates [glucose 6-phosphate (G6P), fructose 6-phosphate (F6P), fructose 1,6-bisphosphte (F1,6BP), and dihydroxyacetone phosphate (DHAP)] were slightly increased by deletion of the *CDC55* gene in the laboratory strain BY4741 (Fig. 5A). In contrast, the level of glyceraldehyde 3-phosphate (G3P) accumulated in *cdc55*Δ cells was 3-fold higher than that in wild-type cells, while the intracellular pools of 3-phosphoglyceric acid (3PG) and the ensuing glycolytic intermediates were smaller in *cdc55*Δ cells. These data suggest that, at 6 h, the metabolic steps between G3P and 3PG are specifically compromised by deletion of the *CDC55* gene. We noted that 1,3-bisphosphoglyceric acid (1,3BPG), an intermediate between G3P and 3PG in the glycolytic pathway, was not detected in both wild-type and *cdc55*Δ cells in the present analysis. At 1 d, similar accumulations were observed for F6P and phosphoenolpyruvic acid (PEP) in *cdc55*Δ cells (Fig. 5B); the accumulation of F6P remained even at 2 d (Fig. 5C). These data suggested that, at 1 - 2 d, the metabolic steps between F6P and F1,6BP, and between PEP and pyruvic acid, are disturbed by deletion of the *CDC55* gene. In the sake strain K701 at 1 d from the onset of alcoholic fermentation, the accumulations of F6P and PEP were observed in *CDC55*^*WT*^-deficient cells (*cdc55*^*WT*^Δ*/cdc55*^*MT*^) (Fig. 5D), consistent with the results obtained using laboratory yeast cells. Based on these results, it is possible to hypothesize that the enzymatic activities of phosphofructokinase (F6P to F1,6BP) and/or pyruvate kinase (PEP to pyruvic acid) are negatively affected by the loss of PP2A^B55δ^ function both in laboratory and sake strains, when the fermentation rates reach their maxima.

**FIG 5.**
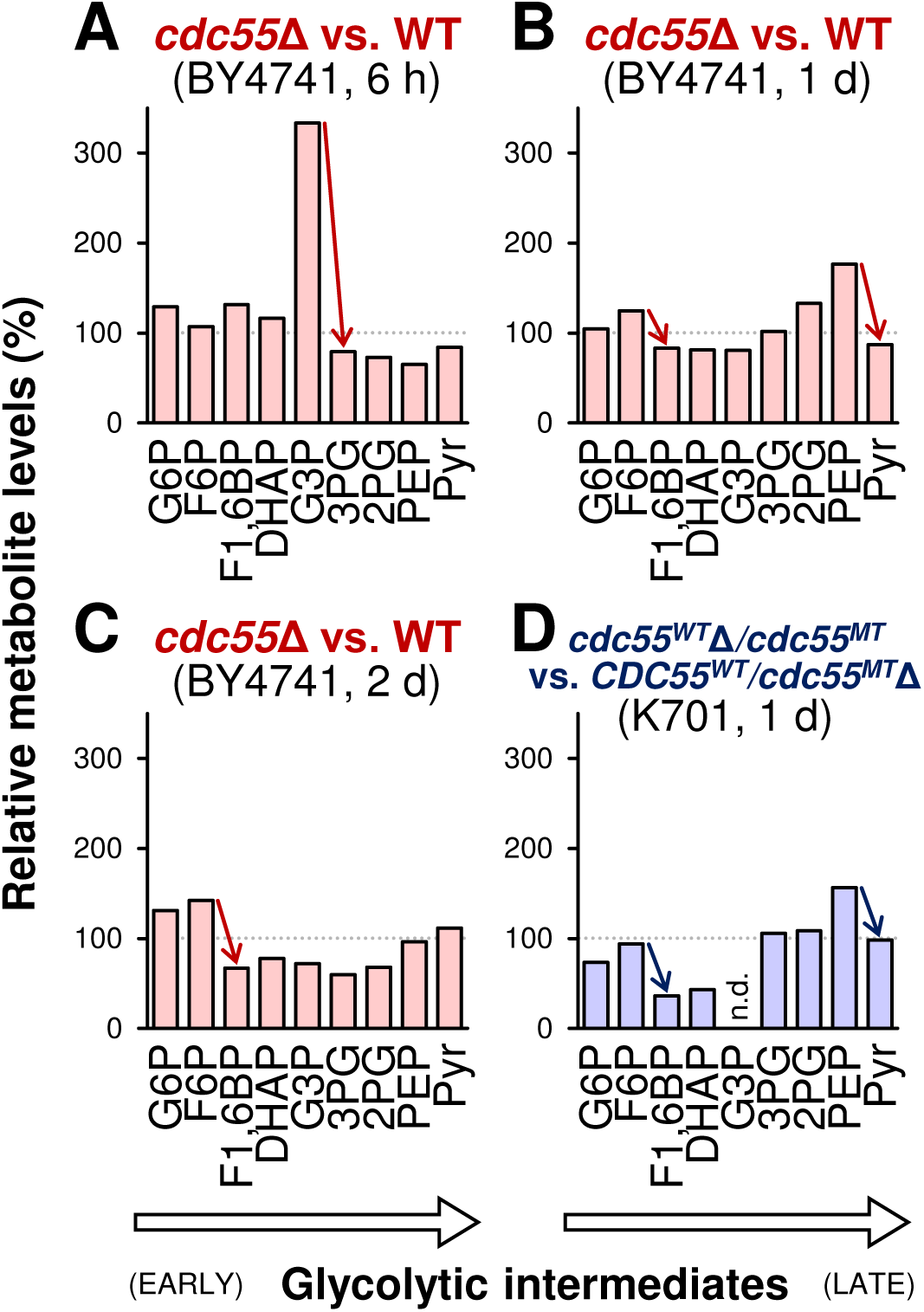
Effects of Cdc55p on glycolytic intermediate levels in the early stage of alcoholic fermentation. (A to C) Intracellular metabolite levels of laboratory strain BY4741 *cdc55*δ at 6 h (A), 1 d (B), and 2 d (C) from the onset of alcoholic fermentation; values are normalized to those of the BY4741 wild type at the respective time point. (D) Intracellular metabolite levels of sake strain K701 *cdc55*^*WT*^Δ/*cdc55*^*MT*^ at 1 d from the onset of alcoholic fermentation; values are normalized to those of K701 *CDC55*^*WT*^*/cdc55*^*MT*^δ. Red and blue arrows indicate notable differences between adjacent metabolites. Data provided are from a single experiment representative of results from multiple independent fermentation tests. G6P, glucose 6-phosphate; F6P, fructose 6-phosphate; F1,6BP, fructose 1,6-bisphosphate; DHAP, dihydroxyacetone phosphate; G3P, glyceraldehyde 3-phosphate; 3PG, 3-phosphoglyceric acid; 2PG, 2-phosphoglyceric acid; PEP, phosphoenolpyruvic acid; Pyr, pyruvic acid; n.d., not determined. Note that G3P was not detected in K701 *cdc55*^*WT*^Δ*/cdc55*^*MT*^, and that 1,3-bisphosphoglyceric acid (1,3BPG) was not detected in any of the samples.

## DISCUSSION

Although the genes and enzymes of the glycolysis and alcoholic fermentation pathways have been thoroughly studied in *S. cerevisiae*, the mechanisms by which intracellular signaling pathways regulate carbohydrate metabolism in response to extracellular cues are still not fully elucidated. We previously identified a loss-of-function mutation in the *RIM15* gene (*rim15*^*5054_5055insA*^) that is present in K7 and shared among the associated sake yeast strains, indicating that this mutation is associated with enhanced fermentation performance (5, 7). In the present work, we showed that sake yeast cells exhibit elevated TORC1 activity during alcoholic fermentation in comparison to laboratory strains. TORC1 upregulates the Sfp1p-targeted genes encoding ribosome-associated proteins and downregulates members of the NCR and GAAC regulons in a Rim15p-independent manner (27). These attributes (Fig. 1), as well as the observed defect in induction of the Msn2/4p-mediated stress-response genes (3), suggest enhanced activation of TORC1 in sake yeast cells (compared to laboratory strains). The high level of phosphorylated Thr737 of Sch9p observed in sake yeast cells (Fig. S2) is consistent with this idea. As previously reported, TORC1 activity is not fully attenuated in K7 cells even under nitrogen limitation (34). Therefore, elevated TORC1 activity can be regarded as a novel hallmark of the sake yeast cells. In general, nutritional limination and environmental stresses rapidly inactivate TORC1 in yeast, resulting in inhibition of cell growth and proliferation. We postulate that the maintenance of high TORC1 activity in sake yeast cells may facilitate cellular metabolic activity even under fermentative conditions. Among the components of TORC1, only Tor1p contains missense mutations (R167Q and T1456I) in K7 and its relatives. Further studies will be needed to evaluate the roles of the mutations in *TOR1* and those in other genes to be discovered in sake yeast strains.

In the present study, we demonstrated that the conserved TORC1-Greatwall-PP2A^B55δ^ pathway is key to the control of alcoholic fermentation (Fig. 6A). In *S. cerevisiae* laboratory strains and *S. pombe*, altered TORC1 activities led to changes in fermentation performance, specifically at the early stage of alcoholic fermentation. However, in laboratory yeast cells deficient for Greatwall, the initial rate of alcoholic fermentation was maintained and not affected by changes in TORC1. In contrast, in PP2A^B55δ^-deficient laboratory yeast cells, the fermentation rate was strikingly low and not enhanced even by a loss of Greatwall or ENSA. The observed strong epistasis suggested that the Greatwall-PP2A^B55δ^ pathway, among numerous downstream effector proteins of TORC1, is the primary mediator of fermentation control. This epistasis also indicated that PP2A^B55δ^ is the major regulator of the alcoholic fermentation machinery.

**FIG 6.**
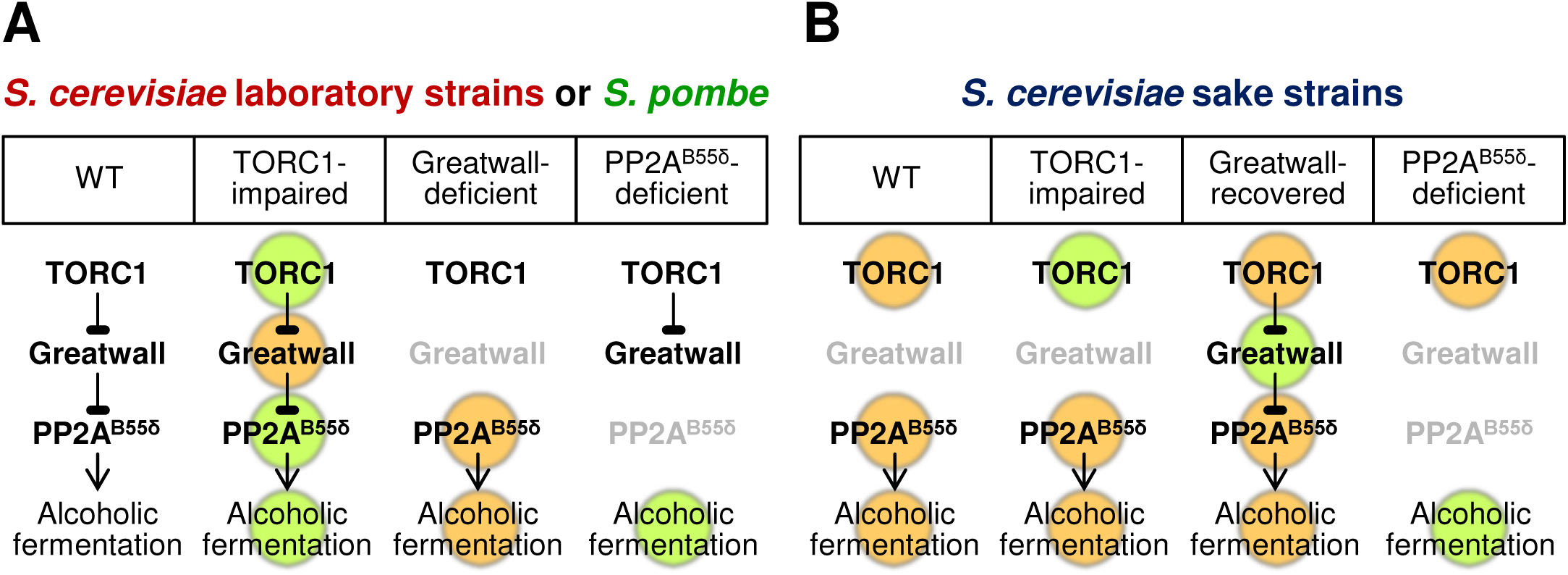
A hypothetical model of the regulation of fermentation control by the TORC1-Greatwall-PP2A^B55δ^ pathway. Orange and green colors indicate higher and lower activities, respectively, than those of *S. cerevisiae* wild-type laboratory strains. (A) In *S. cerevisiae* laboratory strains and *S. pombe*, changes in the activity of TORC1, Greatwall, or PP2A^B55δ^ may lead to altered alcoholic fermentation performance. (B) In *S. cerevisiae* sake strains, both the high TORC1 activity and the loss of Rim15p may contribute to the constitutively high PP2A^B55δ^ activity. Thus, PP2A^B55δ^ must be disrupted to impair the fermentation performance in these strains.

In our hypothesis, both high TORC1 activity and loss of Rim15p contribute to the activation of PP2A^B55δ^ and the subsequent enhancement of the cellular fermentation performance in the K7-related sake strains (Fig. 6B). Indeed, neither impairment of TORC1 nor recovery of Rim15p is sufficient to attenuate PP2A^B55δ^ activity in these cells. Presumably, even if TORC1 activity is decreased, the change in TORC1 signaling may not be conveyed downstream due to the loss of Rim15p (Fig. S1). On the other hand, if a functional *RIM15* gene is restored, the hyperactivated TORC1 can inhibit the functions of Rim15p, resulting in elevated PP2A^B55δ^ activity. Our data indicated that the high fermentation performance of sake yeast cells was abrogated only when the functional *CDC55* (B55Δ-encoding) gene was disrupted (Fig. 3R). Consequently, the two changes (in TORC1 and Rim15p) observed in the TORC1-Greatwall-PP2A^B55δ^ pathway of sake yeast cells may mutually ensure the robust phenotype of these strains in the context of alcoholic fermentation.

Why do multiple sake yeast strains possess putative loss-of-function mutations (i.e., nonsense mutations and frameshift mutations; Fig. 4B) in the *CDC55* gene? Since diploid sake yeast strains contain two copies of the *CDC55* gene, heterozygosity for a loss-of-function mutation at the loci may not yield apparent effects on alcoholic fermentation. PP2A^B55δ^ regulates not only carbohydrate metabolism but also cell cycle progression. In *S. cerevisiae*, PP2A^B55δ^ is the key inhibitor of the entry into quiescence (G_0_ phase). Loss of Rim15p decreases the expression of stress-response genes and shortens chronological life span, and *cdc55*Δ is able to suppress such Rim15p-deficient phenotypes (24). The heterozygous loss-of-function mutations in *CDC55* in the sake strains may reduce the dosage of functional Cdc55p, thereby serving as weak suppressors of the long-term survival defect associated with the *rim15*^*5054_5055insA*^ mutation (Fig. S3). Another mutation in the functional *CDC55* allele or a loss of heterozygosity (LOH) may further enhance cell viability, although the lack of Cdc55p function severely impairs fermentation performance. Thus, the individual sake strains may have independently acquired and maintained the heterozygous *cdc55* mutations during decades of selection for enhanced fermentation. Based on our model, we propose that the *cdc55* mutations identified in the sake strains are potential fermentation inhibitors whose elimination could facilitate the development of genetically stable sake yeast strains.

Comparison of the glycolytic intermediate pools between wild-type and *cdc55*Δ cells suggested that the loss of PP2A^B55δ^ negatively affects the metabolic reactions responsible for the conversion of (i) F6P to F1,6BP, (ii) G3P to 3PG, and (iii) PEP to pyruvic acid during the initial stage of alcoholic fermentation (Fig. 5). We presume that these defects are at least partially responsible for the low fermentation performance of *cdc55*Δ cells. Intriguingly, PP2A^B55δ^ appears to control individual glycolytic reactions in a fermentation-phase-specific manner; only the defect in (ii) was observed at 6 h from the onset of alcoholic fermentation in a laboratory strain, whereas the defects of (i) and/or (iii) were observed from 1 d to 2 d. Thus, these results imply that the activities of glycolytic enzymes are separately regulated during alcoholic fermentation, and that the pleiotropic functions of PP2A^B55δ^ contribute to the optimal glycolytic flux. Among the glycolytic enzymes, phosphofructokinase and pyruvate kinase catalyze irreversible and rate-limiting reactions, (i) and (iii), respectively, in glycolysis. Recent integrated phosphoproteomics data in budding yeast indicate that Pfk1p and Pfk2p (the α and β subunits of phosphofructokinase, respectively) and Cdc19p (the main pyruvate kinase isozyme) form phosphorylation hubs, suggesting that multiple protein kinases phosphorylate these enzymes to modulate their activity, intracellular localization, or protein degradation (32, 33). For example, it has been reported that phosphorylation of residue Ser163 of Pfk2p inhibits the phosphofructokinase activity *in vivo* under gluconeogenic conditions (35). The protein phosphatase activity of PP2A^B55δ^ may directly regulate glycolytic enzymes by counteracting such inhibitory phosphorylation. In fact, Pfk1p and Pfk2p are listed as putative PP2A^B55δ^-dephosphorylated proteins (31). The 3PG kinase Pgk1p, which is involved in reaction (ii), also is a putative PP2A^B55δ^ target. The phosphorylation status and the activities of the candidate enzymes should be compared between wild-type and *cdc55*Δ cells during alcoholic fermentation. Since the glycolytic pathway and the posttranslational modifications of the glycolytic enzymes are often conserved evolutionarily, our study may also offer clues to identify novel key mechanisms of protein phosphorylation-mediated glycolytic control by the TORC1-Greatwall-PP2A^B55δ^ pathway.

## MATERIALS AND METHODS

### Yeast strains

The yeast strains used in this study are listed in Table S1. *Saccharomyces cerevisiae* laboratory strain BY4741 and its single-deletion mutants were obtained from Euroscarf (Germany). Another *S. cerevisiae* laboratory strain X2180 and *Schizosaccharomyces pombe* wild-type strain 972 were obtained from the American Type Culture Collection (ATCC, USA). Sake yeast strains Kyokai no. 7 (K7) and its relatives (K6, K601, K701, K9, K901, K10, K1001, K11, K12, K13, K14, K1401, K1501, K1601, K1701, and K1801) were provided by the Brewing Society of Japan (BSJ, Japan). *S. pombe* strain ED666 *cek1*Δ∷*kanMX* (*h*^*+*^ *ade6-M210 ura4-D18 leu1-32 cek1*Δ∷*kanMX*) was obtained from Bioneer (Korea).

Disruption of the *IGO2* gene in BY4741 *igo1*Δ was performed using a PCR-based method (36) with a gene-specific primer pair and plasmid pFA6a-hphNT (37) as the template to generate BY4741 *igo1*Δ∷*kanMX igo2*Δ∷*hphNT* (*igo1/2*Δ). Disruption of the *CDC55* gene in BY4741 wild type, BY4741 *rim15*Δ and BY4741 *igo1/2*Δ was performed using a PCR-based method (36) with a gene-specific primer pair and plasmid pFA6a-natNT (37) as the template to generate BY4741 *cdc55*Δ∷*natNT* (*cdc55*Δ), BY4741 *cdc55*Δ∷*natNT rim15*Δ∷*kanMX* (*cdc55*Δ *rim15*Δ), and BY4741 *cdc55*Δ∷*natNT igo1*Δ∷*kanMX igo2*Δ∷*hphNT* (*dc55*Δ *igo1/2*Δ), respectively.

The *TOR1*^*L2134M*^ mutation was previously reported as a hyperactive point mutation in the kinase domain of Tor1p (28). Since the mutation site was conserved in the *TOR2* gene, the corresponding mutation was also introduced to generate *TOR2*^*L2138M*^. Disruption of the *RIM15, GTR1, GTR2*, and *SCH9* genes in TM142 wild type or in TM142 *TOR1*^*L2134M*^ was performed using a PCR-based method (36) with a gene-specific primer pair and plasmid pFA6a-kanMX (37) as the template to generate TM142 *rim15*Δ∷*kanMX* (*rim15*Δ), TM142 *TOR1*^*L2134M*^ *rim15*Δ∷*kanMX* (*TOR1*^*L2134M*^ *rim15*Δ), TM142 *gtr1*Δ∷*kanMX* (*gtr1*Δ), TM142 *gtr2*Δ∷*kanMX* (*gtr2*Δ), and TM142 *sch9Δ*∷*kanMX* (*sch9Δ*), respectively.

Heterozygous disruption of the *CDC55* gene in K701 was performed using a PCR-based method (36) with a gene-specific primer pair and plasmid pFA6a-natNT (37) as the template. Correct disruption of the *CDC55*^*WT*^ or *cdc55*^*MT*^ allele was confirmed by genomic PCR and direct DNA sequencing of the PCR product. Homozygous disruption of the *SCH9* gene in IB1401 was performed according to a previous report (32). To overexpress 3HA-tagged Sch9p from a glycolytic gene promoter in IB1401, plasmid p416-3HA-SCH9 (kindly gifted from Prof. Kevin Morano from the University of Texas, USA) was introduced into IB1401.

Disruption of the *ppk18*^*+*^ and *igo1*^*+*^ genes in 972 wild type was performed using a PCR-based method (36) with a gene-specific primer pair and plasmid pFA6a-kanMX (37) as the template to generate 972 *ppk18*Δ∷*kanMX* and 972 *igo1*Δ∷*kanMX* (*igo1*Δ), respectively. The *kanMX* genes in ED666 *cek1*Δ∷*kanMX* and 972 *ppk18*Δ∷*kanMX* were replaced with *natMX* and *hphMX*, respectively, using a one-step marker switch (38) to generate ED666 *cek1*Δ∷*natMX* and 972 *ppk18*Δ∷*hphMX*, respectively. Both strains were mated and sporulated to generate the prototrophic double mutant *cek1*Δ∷*natMX ppk18*Δ∷*hphMX* (*cek1*Δ *ppk18*Δ). To construct the prototrophic mutants *sck1*Δ∷*his7*^*+*^ *sck2*Δ∷*ura4*^*+*^ (*sck1*Δ *sck2*Δ), *ppa1*Δ∷*ura4*^*+*^ (*ppa1*Δ), *ppa2*Δ∷*ura4*^*+*^ (*ppa2*Δ), and *pab1*Δ∷*ura4*^*+*^ (*pab1*Δ), a suitable wild-type strain was mated with JX766 (39), MY1121, MY1122 (40), and MY7214 (41), respectively, and sporulated. The *pab1*Δ and *igo1*Δ strains were mated and sporulated to generate the prototrophic double mutant *pab1*Δ∷*ura4*^*+*^ *igo1*Δ∷*kanMX* (*pab1*Δ *igo1*Δ).

Yeast cells were routinely grown in liquid YPD medium (1% yeast extract, 2% peptone, and 2% glucose) at 30°C, unless stated otherwise.

### Sequencing of the *CDC55* gene

To analyze the *CDC55* sequence, the gene was amplified by PCR with the primer pair CDC55-(−150)-F (5’-GGC AGC TTA ATA CGA TTA CCC C-3’) and CDC55-(+1906)-R (5’-TGG TGA AGT GAT GAA AGA AGT CC-3’), using genomic DNA from the strain of interest as the template. The PCR product was sequenced directly using a BigDye terminator v3.1 cycle sequencing kit (Thermo Fisher Scientific) and primers CDC55-seq2 (5’-TCG AGG TCA AAC TGG AGA GA-3’), CDC55-seq3 (5’-AAA ATC ATT GCT GCC ACC CC-3’), and CDC55-seq4 (5’-TGA TAC CTA TGA AAA CGA TGC GA-3’) on a 3130xl Genetic Analyzer (Applied Biosystems); sequencing was performed at Fasmac Co., Ltd. (Japan).

### Fermentation tests

For measurements of fermentation rates, yeast cells were precultured in YPD medium at 30°C overnight, inoculated into 50 mL of YPD20 medium (1% yeast extract, 2% peptone, and 20% glucose) for *S. cerevisiae* or YPD10 medium (1% yeast extract, 2% peptone, and 10% glucose) for *S. pombe* at a final optical density at a wavelength of 600 nm (OD_600_) of 0.1, and then further incubated at 30°C without shaking. Fermentation progression was continuously monitored by measuring the volume of evolved carbon dioxide gas using a Fermograph II apparatus (Atto) (42).

### Analysis of intracellular metabolite profiles

During the fermentation tests in YPD20 medium, yeast cells corresponding to an OD_600_ of 20 were collected at 6 h, 1 d, or 2 d from the onset of the fermentation tests. All pretreatment procedures for the samples were performed according to the protocols provided by Human Metabolic Technologies, Inc. Briefly, each sample of yeast cells was washed twice with 1 mL ice-cold Milli-Q water, suspended in 1.6 mL methanol containing 5 μM internal standard solution 1 (Human Metabolic Technologies), and then sonicated for 30 s at room temperature. Cationic compounds were measured in the positive mode of CE-TOFMS, and anionic compounds were measured in the positive and negative modes of CE-MS/MS (43). Peaks detected by CE-TOFMS and CE-MS/MS were extracted using automatic integration software (MasterHands, Keio University (44) and MassHunter Quantitative Analysis B.06.00, Agilent Technologies, respectively) to obtain peak information, including *m*/*z*, migration time, and peak area. The peaks were annotated with putative metabolites from the HMT metabolite database (Human Metabolic Technologies) based on their migration times in CE and *m*/*z* values as determined by TOFMS and MS/MS. Metabolite concentrations were calculated by normalizing the peak area of each metabolite with respect to the area of the internal standard and by using standard curves, which were obtained from three-point calibrations.

## ACKNOWLEDGEMENTS

The Japan Society for the Promotion of Science (JSPS) provided funding to DW under grant number 16K18676, to SI under grant number 17H03795, and to TM under grant numbers 25291042 and 17H03802. The Public Foundation of Elizabeth Arnold-Fuji provided funding to DW. The Foundation for the Nara Institute of Science and Technology provided funding to DW. The authors declare no conflicts of interest.

